# Design of Experiments for Fine-Mapping Quantitative Trait Loci in Livestock Populations

**DOI:** 10.1101/2019.12.17.879106

**Authors:** Dörte Wittenburg, Sarah Bonk, Michael Doschoris, Henry Reyer

## Abstract

Single nucleotide polymorphisms (SNPs) which capture a significant impact on a trait can be identified with genome-wide association studies. High linkage disequilibrium (LD) among SNPs makes it difficult to identify causative variants correctly. Thus, often target regions instead of single SNPs are reported. Sample size has not only a crucial impact on the precision of parameter estimates, it also ensures that a desired level of statistical power can be reached. We study the design of experiments for fine-mapping of signals of a quantitative trait locus in such a target region.

A multi-locus model allows to identify causative variants simultaneously, to state their positions more precisely and to account for existing dependencies. Based on the commonly applied SNP-BLUP approach, we determine the z-score statistic for locally testing non-zero SNP effects and investigate its distribution under the alternative hypothesis. This quantity employs the theoretical instead of observed dependence between SNPs; it can be set up as a function of paternal and maternal LD for any given population structure.

We simulated multiple paternal half-sib families and considered a target region of 1 Mbp. A bimodal distribution of estimated sample size was observed, particularly if more than two causative variants were assumed. The median of estimates constituted the final proposal of optimal sample size; it was consistently less than sample size estimated from single-SNP investigations which was used as a baseline approach. The second mode pointed to inflated sample sizes and could be explained by blocks of varying linkage phases leading to negative correlations between SNPs. Optimal sample size increased almost linearly with number of signals to be identified but depended much stronger on the assumption on heritability. For instance, three times as many samples were required if heritability was 0.1 compared to 0.3. These results enable the resource-saving design of future experiments for fine-mapping of candidate variants in structured and unstructured populations.

## 1 Introduction

Genomewide association studies (GWAS) help exploring the relationship between genetic and phenotypic variation. Genetic variation is often expressed in terms of genomic markers such as single nucleotide polymorphisms (SNPs). Identified variants may or may not be part of known genes. In a candidate-gene approach, variants are then assigned to the closest known gene and their functional importance can be studied further (e.g., Reyer *et al*., 2015). The functional meaning of a variant may be differently interpreted if, due to statistical uncertainty, it was identified a few kbp upstream or downstream of its position. In general, it could be a complicated task to detect single loci as reported by Sahana *et al*. (2014) in a study on udder health in dairy cattle. Instead of identifying important SNPs for clinical mastitis, only target regions were found. For instance, a window of about 1Mbp length was detected on BTA6. A statistical reason for this complication lies in the high multicollinearity among predictor variables due to linkage and linkage disequilibrium (LD) between SNPs (e.g., Hampel *et al*., 2018). Region-based aggregation tests in biologically relevant regions (e.g., genes; Lee *et al*., 2014) or fine-mapping approaches in independent partitions of the genome (Schaid *et al*., 2018) have been suggested as powerful options. To eventually unravel which of the variants in a target region might be truly related to the trait, a follow-up experiment is recommended. The experimental design should account for the dependence between SNPs to ensure sufficient statistical power. This will be reflected in the sample size required. Statistical tools for the design of experiments (e.g., QUANTO; Gauderman & Morrison, 2007) could not provide this until now. However, the denser the SNP chip is, the higher will be the correlation between SNPs. For instance, the target region on BTA6 of Sahana’s paper covers 17 SNPs using a 50k SNP panel, 192 SNPs based on a 700k SNP panel and 21 796 SNPs in case of DNA sequence (Sahana *et al*., 2014; Schnabel, 2018).

In theory, it can be determined what sample size is needed for discovering a new variant in a single-locus model at a given power, e.g., 80 %. Such investigations are based on ANOVA (one way classification; Luo, 1998) and can also account for a hypothetical degree of LD between causative variant and SNP (Pritchard & Przeworski, 2001; Khatkar *et al*., 2008). Proposals for an optimum experimental design have been made for mapping of a quantitative trait locus (QTL) in different population structures (e.g., F2, backcross or daughter design; Weller, 2001). However, it is not clear what sample size is required to distinguish multiple independent signals of a QTL using dense marker data.

Moreover, the power of association analysis depends not only on sample size and population parameters (e.g., heritability) but also on the underlying statistical model. Among myriad options for whole genome regression models, SNP-BLUP is an obvious choice for estimating genetic effects captured by all SNPs simultaneously. Also because of its direct relationship to GBLUP (e.g., Gualdrón Duarte *et al*., 2014), it is widely used in livestock (e.g., Koivula *et al*., 2012; Mucha *et al*., 2015) and beyond (e.g., Maier *et al*., 2015; Kristensen *et al*., 2018). Being enormously relevant in practice, it has been upgraded to comprise information of individuals with and without genotypic data in the framework of single-step methods (Taskinen *et al*., 2017; Aguilar *et al*., 2019). Though directly or indirectly estimated SNP effects are tested for being significantly different from zero (Aguilar *et al*., 2019), reports on statistical power of the underlying study design are lacking.

This paper addresses the question how to design a follow-up experiment based on a SNP-BLUP approach knowing that the predictor variables are so highly correlated. Our objective is the theoretical inference of optimal sample size to fine-map a QTL signal or to find evidence for multiple independent signals in a specified chunk of DNA. Eventually, it should be possible to detect variants at their actual position with high power. This paper concentrates on the case study of paternal half-sib families which is a typical family structure in livestock (e.g., dairy cattle). But the methodology developed enables sample size calculation for any population structure (e.g. full siblings, half siblings, mixture of both, unrelated individuals). Given the number of families, SNPs, signals of QTL and heritability, the optimal sample size is then presented as overall number of progeny. We validated our approach using simulated data. Furthermore, publicly available bovine HD SNP chip data helped verifying that the simulated linkage blocks resemble the genome structure in dairy cattle. A discussion of our achievements complements this study.

## 2 Material and methods

The design of experiment requires a statistical model that combines phenotype with genotype data. Here, we assume a multiple-SNP approach that considers information of as many SNPs as desired simultaneously. For comparing the outcome with a conventionally used approach, a single-SNP model is specified.

### 2.1 Multi-SNPs model

For a joint association analysis of *p* SNPs, a regression model is fitted to a phenotype *y* = (*y*_1_, …, *y*_*n*_)′ of *n* individuals,

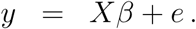

The *n* × *p* design matrix *X* contains the genotype codes: *X*_*j,k*_ ∈ {1, 0, −1} for *j* = 1, …, *n* and *k* = 1, 2, …, *p*. The columns of *X* and the vector *y* are centered within family and scaled afterwards. This way, the model becomes independent of allele frequency. The residual error term is 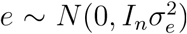. Then the coefficient vector *β* is estimated using a ridge regression approach as

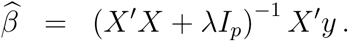

This step requires a penalty term λ which is practically obtained via cross-validation or REML approach.

Next, we investigate a multiple testing problem. For each SNP *k, k* = 1, …, *p*, it is tested

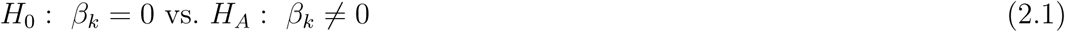

with a suitable test statistic

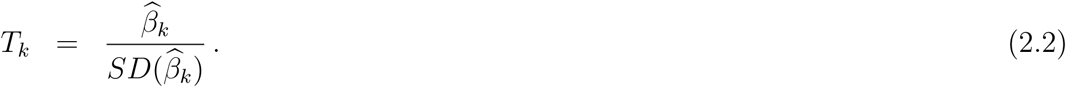

The calculation of power *π* requires the distribution of *T*_*k*_ under *H*_*A*_, then

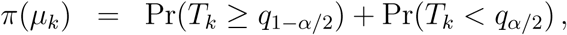

where *q*_1−*α/*2_ and *q*_*α/*2_ denote the upper and lower threshold, respectively, of the distribution of *T*_*k*_ under *H*_0_ with respect to a type-I error *α*. Due to the ridge approach, requirements for fulfilling a *t* distribution do not hold (Searle, 1971, p. 57). Hence the distribution of *T*_*k*_ is approximated as normal and the distribution mean *µ*_*k*_ is obtained from the expectation and variance of the estimator 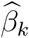. The moments are

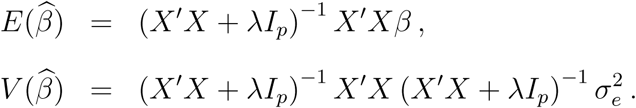

The central point of our investigation is to substitute the correlation matrix 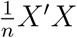 to be observed in the progeny generation by the theoretical correlation matrix *R*. The calculation of *R* requires a genetic map and genetic information of parents; its derivation is shown in Appendix A.

Then the mean becomes

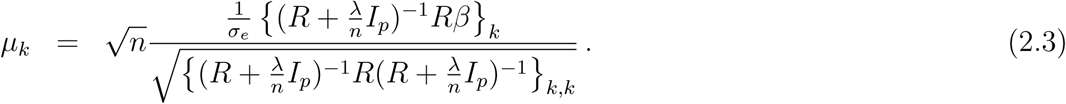

Under *H*_0_ *µ*_*k*_ = 0. In order to calculate the optimal sample size, the experimenter has to specify a set of parameters: number of SNPs (*p*) in the investigated window of DNA, number of QTL signals to be detected (*η*), proportion of variance explained by the window (*h*^2^) and number of families (e.g., *N* sires). The input parameters for statistical power calculation are inferred from this experimental set-up:

1. *R* requires haplotypes of *N* sires (plus genetic map and maternal LD in general).
2. We assume that all variants corresponding to the QTL signals contribute equally to the genetic variance. Hence the relative effect size is determined at *η* QTL signals as

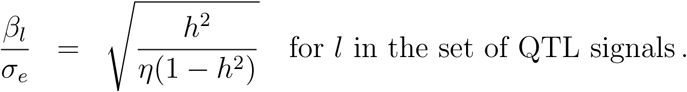 The remaining *β*’s are 0.
3. The shrinkage parameter is derived corresponding to Hoerl *et al*. (1975),

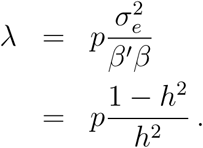

This is a rough approximation assuming linkage equilibrium between variants corresponding to the QTL signals.

We circumvent doing any assumption about the unknown positions of QTL signals by taking a random sample of *η* positions. Then the optimal sample size is calculated over a range of *n* (e.g., 1 − 5 000) employing the method of bisection. The minimum *n* that exceeds a power of 80% is selected as “optimal” and denoted as *n*_opt_. In order to get a reliable estimate of optimal sample size, sampling is repeated 100 times, and the median of *n*_opt_ is suggested as final 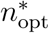. The overall type-I error was *α* = 0.01.

### 2.2 Single-SNP model

For comparison, we consider a single SNP *k* ∈ {1, …, *p*} in a sliding window over the target region. Using the parameter definitions as above, the linear model in its simplest form is

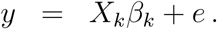

Then the regression coefficient is estimated via ordinary least squares as

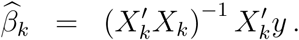

The null hypothesis testing problem (2.1) and the corresponding test statistic (2.2) also apply in the single-SNP analysis. The test statistic is *t*-distributed with *n* − 1 degrees of freedom and non-centrality parameter *δ*_*k*_ (Searle, 1971, pp. 110),

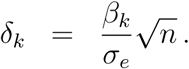

This approach neglects any impact of the other SNPs in the target region on *y*. Thus, a reduced pointwise error level (*α*_*k*_) has to be employed to keep the overall type-I error at *α*. Knowing the effective number of independent tests (*M*_eff_), a suitable type-I-error correction is

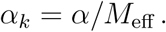

In accordance with the simpleℳ method of Gao *et al*. (2008), we suggest using *R* instead of 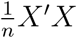 for calculating *M*_eff_. More precisely, the number of eigenvalues of *R* that contribute at least 99.5% to the sum of all eigenvalues yields *M*_eff_.

### 2.3 Data and validation study

The software R version 3.6.1 (R Core Team, 2019) was employed in this study. Unless otherwise stated, we implemented own R scripts.

Population genetic data were simulated using the R package `AlphaSimR` version 0.11.0 (Faux *et al*., 2016). In total, 300 SNPs were spread in a chunk of DNA of 1cM length. This corresponds approximately to 1Mbp. Five traits were simulated simultaneously, one for each number of QTL signals affecting the trait, *η* = 1, …, 5. In case of multiple QTL signals, effect sizes were equal. The founder population comprised 2 000 individuals and constitutes the parent generation. Other population parameters were kept at default settings. As the data simulation yielded no consistent pattern of SNP dependence, the simulation of the parent population was repeated 100 times. The maternal LD in terms of *r*^2^ between adjacent SNPs was on average 0.45 and reflects high multicollinearity. In each repetition, *N* = 10 sires were selected depending on their phenotype. Then, haplotypes of all selected sires, or a subset thereof if *N* = 1 or *N* = 5, and 1 000 females were used to set up the *R* matrix. Few loci with no variation were disregarded. Optimal sample size was estimated based on *R*. Using the same parent generation, *N* males were selected as sires of half-siblings in the progeny generation. The number of dams was determined according to optimal sample size required. The simulation of the progeny generation was also repeated 100 times to estimate and test SNP effects for validation purposes; this yielded 100 × 100 data sets in total. The total heritability was *h*^2^ ∈ {0.1, 0.2, 0.3}. For each *h*^2^, data sets were simulated independently.

Additionally, to explore a direct relationship between assumed positions of QTL signals and *n*_opt_, we selected arbitrarily a single repetition of simulation with *h*^2^ = 0.1 and *N* = 10. For this particular data set, we determined *n*_opt_ for each SNP position (i.e., assuming one QTL signal) and for all possible SNP pairs (i.e., assuming two QTL signals).

The R package `asreml` version 3.0 (Butler *et al*., 2009) was used for association analysis. Other suitable R packages, such as `rrBLUP` (Endelman, 2011) or `ridge` (Cule *et al*., 2011), had difficulties to converge or produced almost zero variance components due to the high multicollinearity of predictor variables. The multi-SNP model was applied to all simulated scenarios as described in Section 2.1. Unlike in Section 2.2, the single-SNP model considered an additional factor 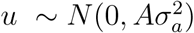 that accounts for background genetic effects due to the relationship between individuals. This was modeled similarly, e.g., in EMMAX (Kang *et al*., 2010) but we used the numerator relationship matrix *A* for computational convenience. The pointwise testing of SNP effects was followed by *p*-value correction according to Benjamini & Hochberg (1995). *P*-values from the multi-SNP model were not altered. The outcome was used to assess sensitivity and specificity of the multi-SNP and single-SNP model. For this, a window of 0.01 cM to both sides of a QTL signal (covering 2-3 SNPs) was specified in order to accept a significant SNP as a true positive result. Then, the true-positive rate (TPR) reflected sensitivity. Specificity was obtained as 1− the false-positive rate (FPR), and ROC curves were produced from TPR and FPR.

To evaluate how realistically the simulation of genetic data worked, empirical HD SNP chip data from the Dryad repository have been used (Bermingham *et al*., 2013). These data included 1 151 dairy cows with no pedigree specification. We selected an arbitrary window on BTA7 comprising 300 SNPs on 1.16 Mbp and phased haplotypes of all animals using AlphaPhase (Hickey *et al*., 2011). We selected randomly 10 animals and marked them as sires in order to set up a matrix *R*. Because of the high SNP density, genetic distances were approximated linearly, i.e., 1Mbp ≈ 1 cM. Maternal LD was roughly approximated from haplotype frequencies of all animals.

### 2.4 Data availability

R scripts used to simulate and analyze data are available upon request. The empirical bovine HD SNP chip data are accessible through the Dryad repository https://doi.org/10.5061/dryad.519bm (Bermingham *et al*., 2013).

## 3 Results

The optimal sample size suggested by the single-SNP model required the effective number of independent tests which was on average *M*_eff_ = 53 if *h*^2^ = 0.1 and rather constant for *R* set up from *N* = 1,5 or 10 sires (*h*^2^ = 0.2: *M*_eff_ = 54; *h*^2^ = 0.3: *M*_eff_ = 56). Hence results are reported for *M*_eff_ based on *N* = 10. Table 1 presents the median of 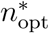 from 100 repetitions of simulation. The median increased almost linearly with number of QTL signals but reduced with increasing heritability, and it was rather unaffected by the number of families. As an example, 127 individuals were required to fine-map a single QTL signal based on the multi-SNP model if *h*^2^ = 0.1. Almost twice as much were required to distinguish two signals if *h*^2^ = 0.1 or only 34 individuals were required to detect a single signal correctly when *h*^2^ = 0.3 instead of *h*^2^ = 0.1. Optimal sample size suggested by the multi-SNP model was 17% to 39% less than estimated from the single-SNP model. Supplemental Figure D.1 visualizes the dependence of optimal sample size estimated from the single-SNP model on heritability. It also shows that a much larger sample was required if QTL heritability was less than 0.2.

**Table 1.**
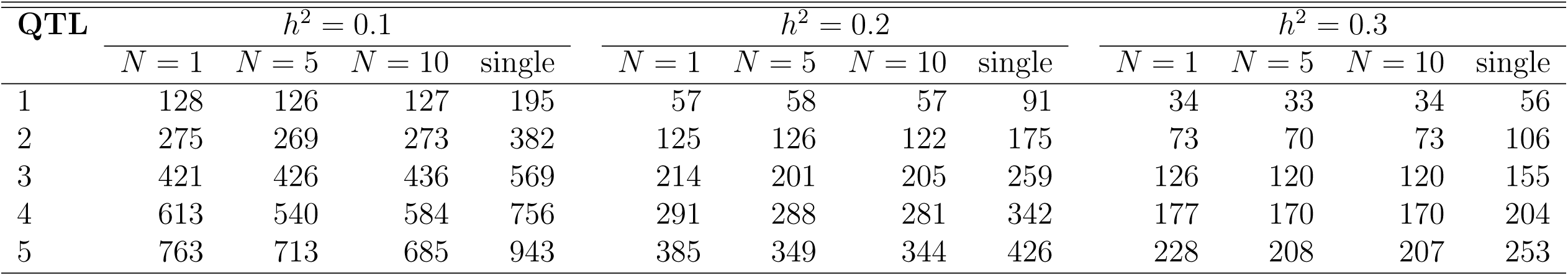
Median of optimal sample size for detecting different number of QTL signals from 100 repetitions of simulations. Results are based on the multi-SNP approach* (*N* = 1, 5, 10 families) or single-SNP approach. (*In each repetition, sample size was repeatedly determined for randomly drawn QTL positions and the median was calculated.)

In case of *h*^2^ = 0.1, the distribution of *n*_opt_ is represented in Figure 1; a separate panel is shown for each number of QTL signals to be detected. Based on 100 × 100 estimates of *n*_opt_, we derived a bimodal distribution of optimal sample size in the multi-SNP model. The median of *n*_opt_ was consistently less than sample size estimated from single-SNP investigations. With increasing heritability, the first mode approached the median of *n*_opt_ but was still less than optimal sample size based on the single-SNP model, see Supplemental Figures D.2 (*h*^2^ = 0.2) and D.3 (*h*^2^ = 0.3). The second mode appeared due to strong negative correlations between SNPs. Particularly this outcome was observed when all possible pairs of SNPs were evaluated for detecting two QTL signals in a single repetition of simulation. Figure 2(a) shows the correlation matrix for a single data set. Those entries of *R* have been selected that belonged to 10% of the highest estimates of sample size, i.e., *n*_opt_ ≥ 864 (*h*^2^ = 0.1). Correspondingly, Figure 2(b) indicates that, with few exceptions, negative correlations caused this outcome. The separation of SNP dependence into maternal and paternal contribution revealed further insight, and most often negative maternal LD was the driving term (Supplemental Fig. D.4). An additional inspection of any relationship between assumed positions of a single QTL signal and *n*_opt_ was not conclusive. Neither extreme maternal allele frequency nor missing sire heterozygosity led to obviously increased *n*_opt_ for detecting one QTL signal (Figure 3).

**Figure 1.**
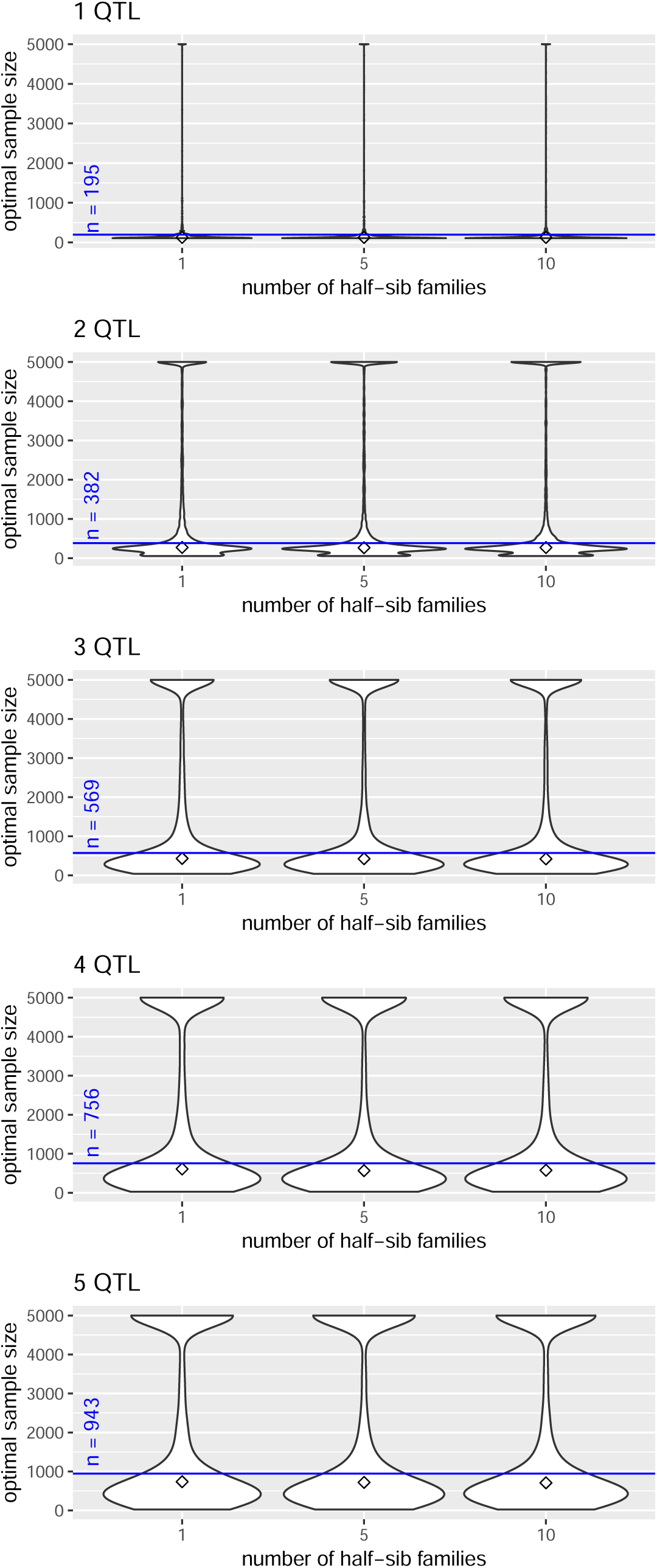
Distribution of optimal sample size. Violinplot of *n*_opt_ vs. number of half-sib families for different numbers of QTL signals in a multi-SNP model. The parent generation was simulated 100 times and 100 random draws of positions of QTL signals were analyzed in each run, *h*^2^ = 0.1. The diamond indicates the median of *n*_opt_ and the blue line marks the results based on a single-SNP model.

**Figure 2.**
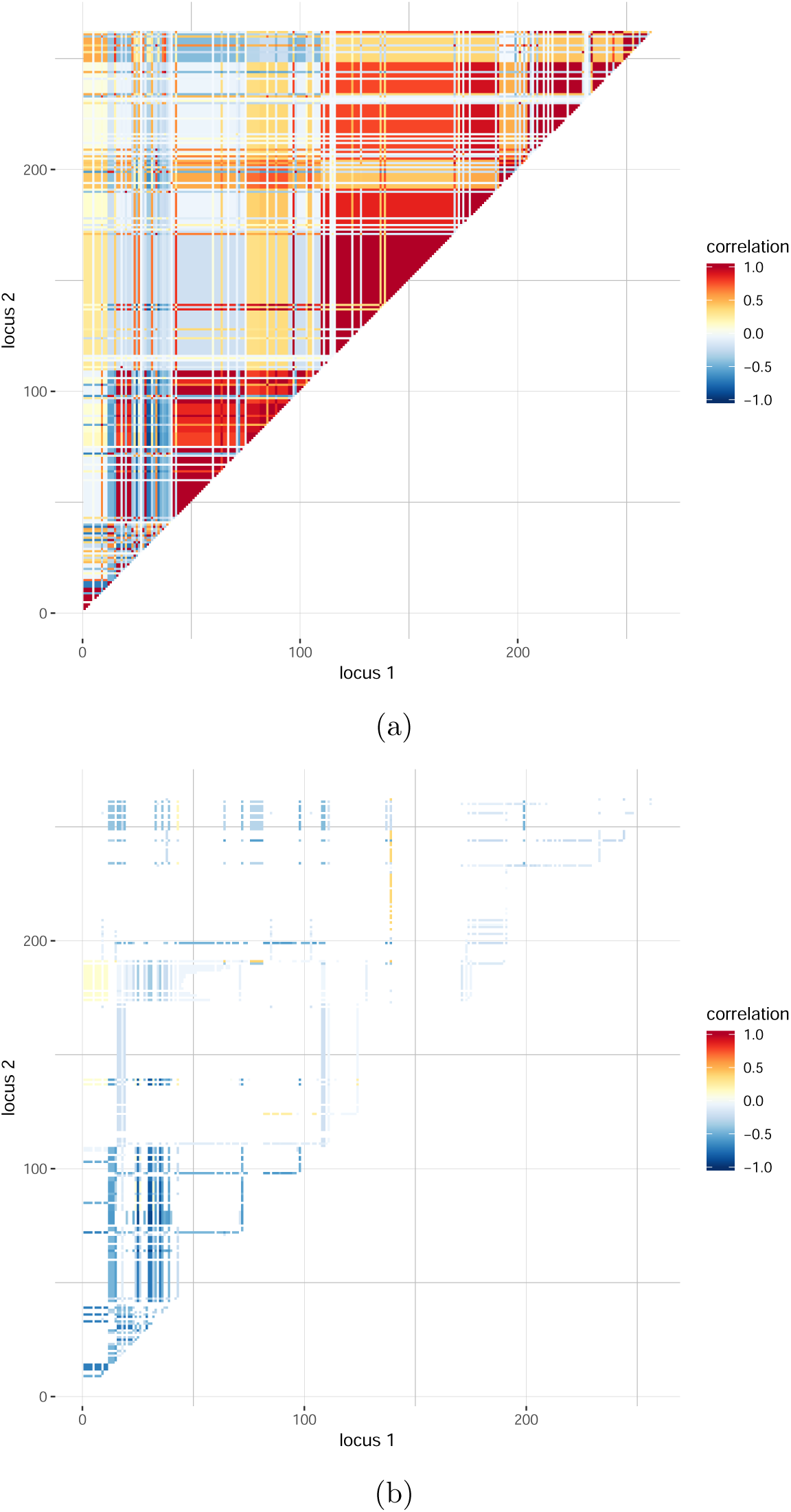
Dependence between SNPs in a single simulated data set with *N* = 10 sires. (a) Correlation matrix *R*, (b) entries selected from *R* which belong to 10% highest sample size (*n*_opt_ ≥ 864). All possible SNP pairs were evaluated to detect two QTL signals (*h*^2^ = 0.1).

**Figure 3.**
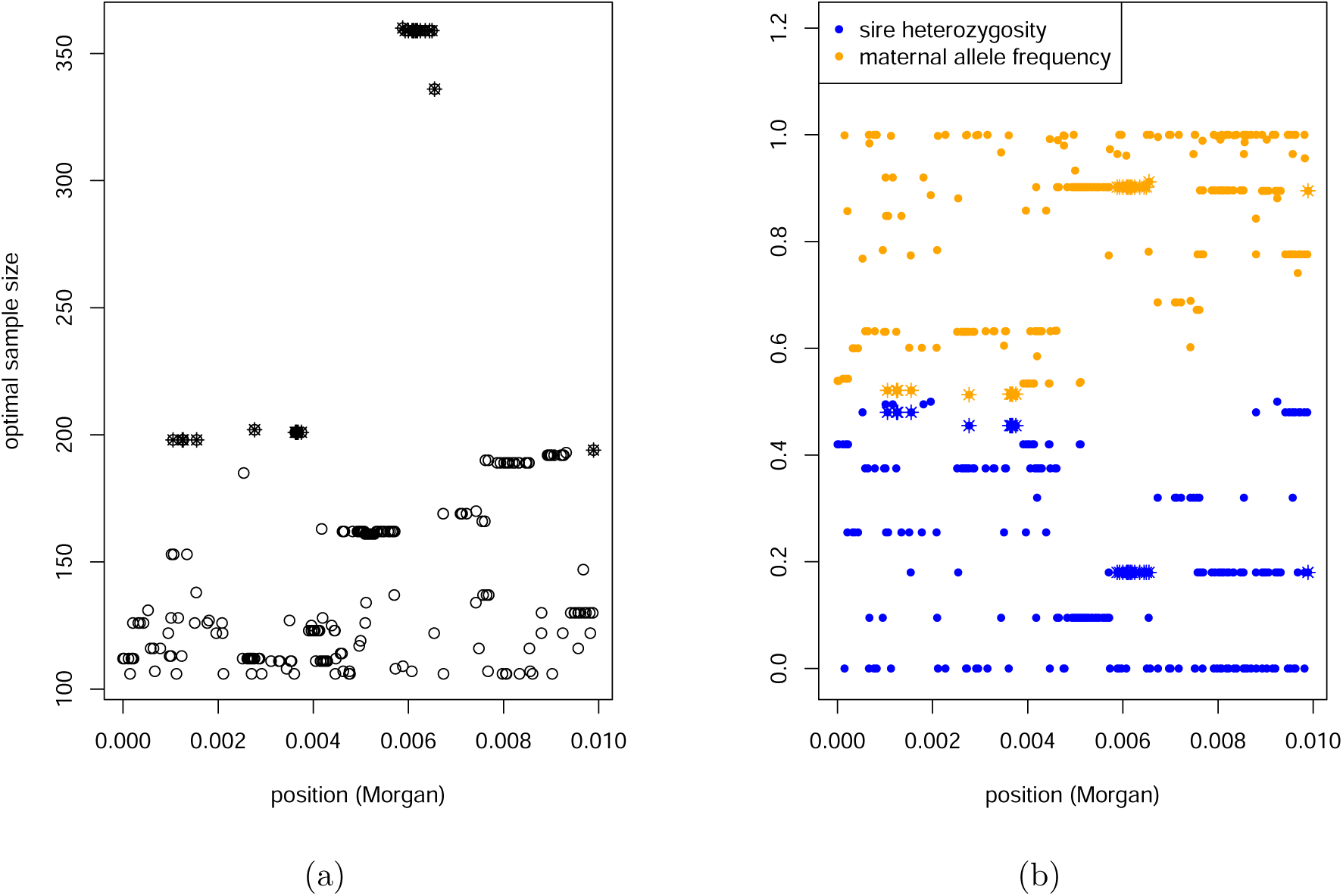
Relationship of optimal sample size with genome position. (a) Optimal sample size for detecting one QTL signal was estimated based on the multi-SNP model (*h*^2^ = 0.1). All possible SNP positions were evaluated. (b) Sire heterozygosity and maternal allele frequency at each SNP position. Values for SNPs that belong to 10% highest sample size (*n*_opt_ ≥ 194) are indicated by a star. Results are based on a single simulated data set with *N* = 10 sires.

The association analysis of data sets of optimal sample size was validated in terms of sensitivity and specificity of testing SNP effects. The shape of ROC curves was similar for all investigated simulation scenarios. As an example, if *N* = 10 and *η* = 2, the median of 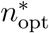 was 273, and the outcome is displayed in Figure 4. The analysis showed superiority of the multi-SNP model over the single-SNP model. In general, it was observed that the smaller 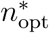 was estimated, the larger both TPR and FPR turned out for the single-SNP model. The multi-SNP model performed rather robust against changes in sample size. However, the flat appearance of the ROC curve complicates fine-mapping of QTL signals based on the suggested multi-SNP approach. For instance, a TPR of 80% is accompanied with a FPR larger than 20 %.

**Figure 4.**
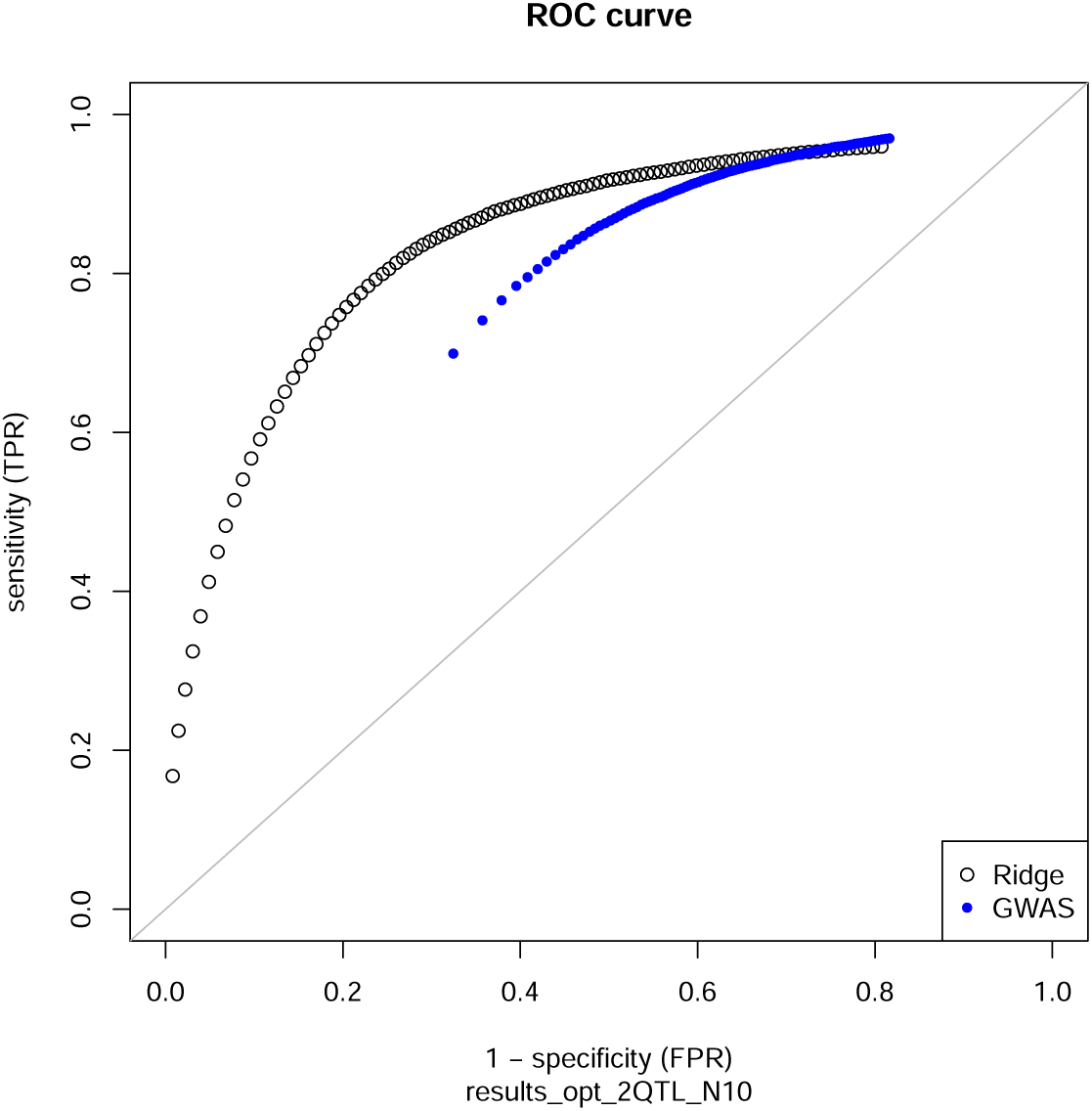
Sensitivity and specificity of testing SNP effects. ROC curve is based on 100 × 100 repeated simulations of genotypes and phenotypes in progeny generation comprising *N* = 10 half-sib families (two QTL signals, *h*^2^ = 0.1). Optimal sample size suggested by the multi-SNP model was considered for setting up the progeny generation.

Blocks of varying linkage phases, as shown in Figure 2, might be an artifact of data simulation. Based on empirical bovine HD SNP chip data, a possible *R* matrix was set up, see Figure 5. The blocking structure was less pronounced. Using this *R* for estimating 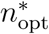 led to results being similar to the simulation study for one and two QTL signals but larger samples were required to detect more QTL signals: 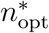 was 123 (1 signal), 288 (2 signals), 516 (3 signals), 800 (4 signals) and 1 342 (5 signals) if *h*^2^ = 0.1. The number of repetitions of randomly drawing the positions of QTL signals did not substantially affect the final 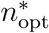. For instance, the median deviated less than 4% if *n*_opt_ was calculated 1 000 instead of 100 times.

**Figure 5.**
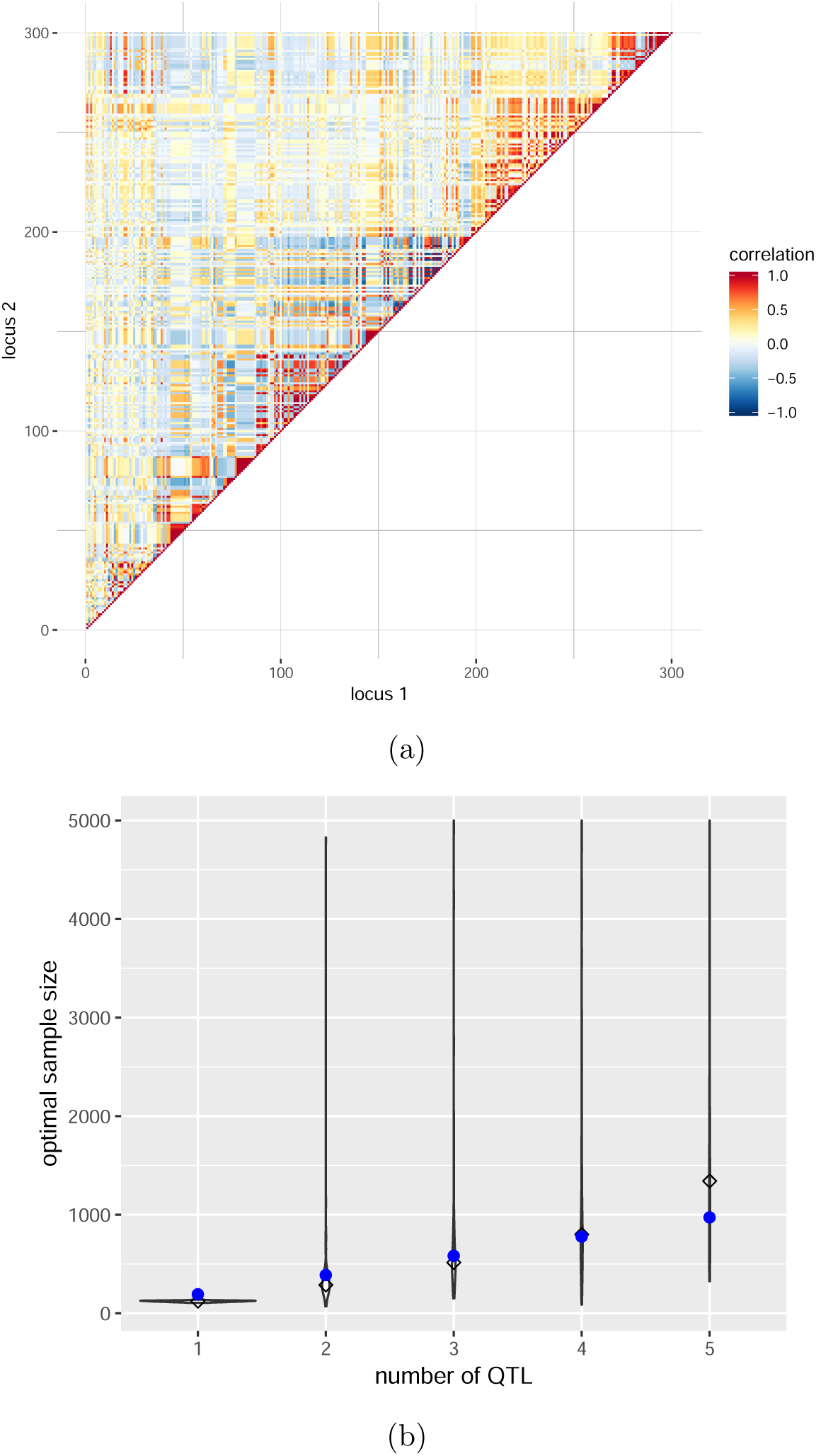
Empirical bovine HD SNP chip data. (a) Correlation matrix for a randomly selected window containing 300 SNPs on BTA7. (b) Violinplot of *n*_opt_ vs. number of QTL signals to be detected. The diamond indicates the median of *n*_opt_ and the blue dots mark the results based on a single-SNP model, *N* = 10 and *h*^2^ = 0.1.

## 4 Discussion

Our investigation contributes to the design of powerful experiments for fine-mapping of causative variant(s) in a genomic target region. We incorporated the expected dependence among SNPs in this region and estimated optimal sample size based on a SNP-BLUP approach. The outcome was compared to a single-SNP model. Negative correlations between SNPs, which were mainly due to negative maternal LD, caused essentially inflated sample size estimates. In case of positive correlations, the majority of sample size estimates was less than sample size estimated from the single-SNP approach. The less the heritability, the higher the deviation between models was.

### 4.1 Population parameters

Our approach is applicable to any population structure. The matrix *K* of covariance between SNPs can be set up for any kind of family stratification by adapting the derivations of the Appendix or, in case of unrelated individuals, by using population LD in *K*.

Due to the way of model parametrization (columns *X*_*k*_ have been scaled), the dependence on allele frequency has been excluded. For instance, in a random mating population, the column-scaling term is 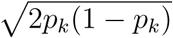 with allele frequency *p*_*k*_ at SNP *k*. Likewise, a scaling term can be derived for half-sib families as the square root of Equation (A.3) by investigating maternally and paternally inherited SNP alleles separately. Results of association analyses suggested that there was no clear relationship between high *n*_opt_ and maternal allele frequency or sire heterozygosity (Figure 3). However, regions with large or low variation have to be taken into account when selecting sires for fine-mapping of QTL signals in a follow-up experiment. The lower sire heterozygosity or maternal minor allele frequency is, the lower the relative effect size *β*_*k*_ on the model scale will be and, consequently, higher 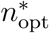 is required in order to detect QTL signals. To investigate this, we employed the relationship 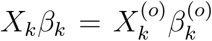 at SNP *k*. Here 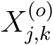 is the allele count at SNP *k* for individual *j*, and 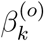 is the coefficient on the observed genotype scale, i.e. 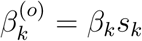 with scaling term 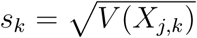. The relationship between allele frequency and optimal sample size for detecting one QTL signal based on the single-SNP model is presented in Supplemental Figure D.5. Extreme alleles, roughly spoken with major allele frequency > 0.95, require drastically increased sample size.

Hence the suggested optimal sample size is divided into *N* sires which are selected for most heterogeneity in the target region. The actual number of sires is of minor importance. The choice of individuals depends on the objectives of the follow-up study. Sires can be chosen independently from the GWAS population in order to confirm and fine-map QTL signal(s). However, if the initial study indicated the presence of rare variants, sires under suspicion should be re-used. Selective genotyping is an option to increase power (Weller, 2001) but this might have negative impact on reproducibility of the study design (Lee *et al*., 2014). In our investigation of paternal half-sib families, mothers are treated as random samples from a dam population. Thus, the choice of dams for future matings is not addressed here but is definitely an issue for other family designs.

Being equally important for fine-mapping of QTL signals is the positive correlation between SNPs. Positive correlatedness is a matter of genotype coding. Coding has to be consistent throughout the target region to avoid unnecessary sign changes in correlation. We employed coding in terms of counting the major allele in the population. But in regions of intermediate frequency, the coding might not be appropriate and hence a dynamic approach of coding the SNP alleles can circumvent negative correlations. A strategy on this is worth further investigation.

Power calculations are needed to quantify and judge the prospects of identifying causative variants with a hypothesized effect size in a particular population. In practice, however, experiments are usually not planned to obtain maximum power but data are regularly collected for purposes of breeding as a standard routine. Thus, experimental designs being theoretically optimal could be compared with available field data to understand the possible shortcomings of such data and to understand differences between theoretical/expected and actually achieved power. Based on the results, decisions can be made whether the amount of data is sufficient or, in case of underpowered experiments, more data should be acquired.

### 4.2 Necessity of fine-mapping of QTL signals using an appropriate design

The QTL databases of livestock species (Hu *et al*., 2012) contain information on several thousands QTL for a wide range of traits. This shows that the variability of most of the traits studied has a polygenic origin, with multiple QTL contributing to the overall genetic variance. Despite the number of QTL, only a handful of causal mutations could be detected and verified in the different livestock species (Andersson & Georges, 2004). This is partly due to the fact that GWAS show considerable weaknesses in the fine-mapping of QTL signals which are related to the SNP panel requirements for a genomewide distribution and high LD to neighboring markers (Schaid *et al*., 2018). Accordingly, these SNPs are usually indicative of a large genomic region that likely comprises the unmeasured causal SNP but does not provide information about the causal variant itself. Statistical methods for fine-mapping have been designed to overcome these issues and perform fine-mapping using the available SNP information from a SNP-chip or GBS (summarized by Schaid *et al*., 2018). However, even these methods require a high SNP density in the region of interest, which favors a targeted sequencing strategy that enable the dissection of QTL regions and increase the chance of detecting causal variants (Mamanova *et al*., 2010). Major factors to be considered for designing a targeted sequencing study are effect size, the number of causal SNPs, local LD structure and sample size (Schaid *et al*., 2018). The approach proposed in this study incorporates information on *η, h*^2^ and *R* derived from the data to estimate the optimal sample size 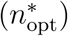 and thus provides all the information needed to design a fine-mapping experiment.

Currently, several fine-mapping studies are based on imputation strategies or the integration of results with functional enrichment analysis to identify promising candidate genes and QTNs (e.g., Jiang *et al*., 2019; Cai *et al*., 2019; Liu *et al*., 2019). These approaches largely depend on imputation accuracy and the status of genome annotation, thus limiting the ability to detect causal variants, especially those with a low allele frequency (Dadaev *et al*., 2018). Specific examples for the fine-mapping of important genomic regions with a resequencing strategy are still rare nowadays. Fraser *et al*. (2018) focused in their study on collagenous lectins in horses by resequencing 658kb DNA consisting of different candidate genes and regulatory regions. Therefore, a case-control design with pooled samples was used and with this approach 113 variants were identified, which differed between the groups. Although the results are promising, the authors concluded that a large-scale genotyping of individual samples is necessary for deeper insights. In this context, and considering that targeted sequencing for a reasonable set of samples is becoming increasingly affordable, an accurate estimate of sample size is advisable.

### 4.3 Other random effects

Association analysis of empirical data with certain pedigree structure requires an additional model term to account for genetic effects beyond the target window (*Zu*). Then 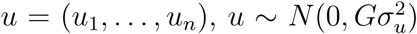, is the vector of individual genetic effects with suitable relationship matrix *G*. The calculation of optimal sample size should consider the presence of additional random effects (genetic or environmental) for the design of experiments. For instance, the coefficients of the single-SNP model could be estimated via BLUE. This affects sample size calculation because the variance of the estimator 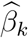,

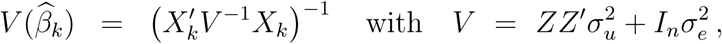

has impact on the distribution of the test statistic. Accounting for *V* in the denominator of test statistic increases the denominator of non-centrality parameter. However, in order to keep it simple, it would be sufficient to increase 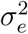 or reduce *h*^2^ appropriately without any other alterations.

It is possible to consider other kinds of genetic effects with the proposed methods. For instance, exploring dominance genetic effects requires only one modification. Instead of coding SNP genotypes for additive effects via *X*_*j,k*_ ∈ {1, 0, −1}, genotypes can be coded as *X*_*j,k*_ ∈ {0, 1, 0} to account for dominance effects. The covariance between dominance effects has been worked out by Bonk *et al*. (2016). Feeding the Equation (2.3) with the corresponding dominance correlation matrix will provide estimates of optimal sample size to fine-map QTL signals with dominance effect.

## 5 Conclusion

For planning the design of experiment, we recommend a multi-SNP approach which considers the expected dependence among SNPs. Compared to a conventional approach, this leads to a reduced estimate of sample size and thus promises a more efficient use of animal resources. The benefit depends strongly on heritability: the lower heritability, the more resources can be saved. In general, optimal sample size increases almost linearly with the number of QTL signals to be detected. This study constitutes a framework for the design of experiments in specific populations that may be characterized by family stratification. It will help differentiating independent signals in QTL regions that can be further examined for cellular and molecular properties.

## Acknowledgments

The project was funded by the German Research foundation (DFG, WI 4450/1-1). Special thanks are given to N. Reinsch (Leibniz Institute for Farm Animal Biology, Dummerstorf), J. Hartung (University of Hohenheim, Stuttgart) and V. Liebscher (University of Greifswald), who contributed invaluable ideas to the project, and to C. Gaynor (Roslin Institute, Edingburgh) for giving handy insight into AlphaSimR.

## Authors’ contribution

DW developed the theory, implemented the statistical methods, performed the analysis, and wrote the manuscript. SB contributed to the research on covariance between SNPs, MD was involved in theoretical investigations. HR contributed to the discussion. All authors have read and approved the final manuscript. The authors declare that they have no competing interests.

## A Appendix Derivation of correlation matrix

We study the dependence between pairs of SNPs, each with two alleles A and B, in a population consisting of *N* paternal half-sib families. Let *X*_*j,k*_ be the genotype code at SNP *k* ∈ {1, …, *p*} of individual *j* ∈ {1, …, *n*} being progeny of sire *s* and dam *d*. Homozygous genotypes A/A and B/B are coded as 1 and -1, respectively, and the heterozygous genotype A/B is indicated as 0. The family-specific, i.e. sire-specific, covariance between SNP *k* and *l* of individual *j* is, according to Bonk *et al*. (2016) and Wittenburg *et al*. (2016),

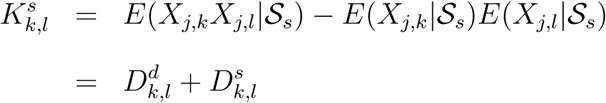

a function of maternal and paternal contribution and depends on the sire diplotype 𝒮_*s*_. The 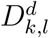 denotes the LD of maternal gametes in a dam population. The sire term depends on the phase of paternal haplotypes and recombination rate (*θ*_*k,l*_). It is determined as

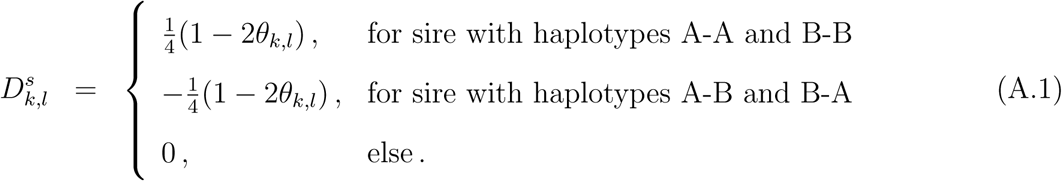

To achieve the covariance between a pair of SNPs, we employ conditioning on families,

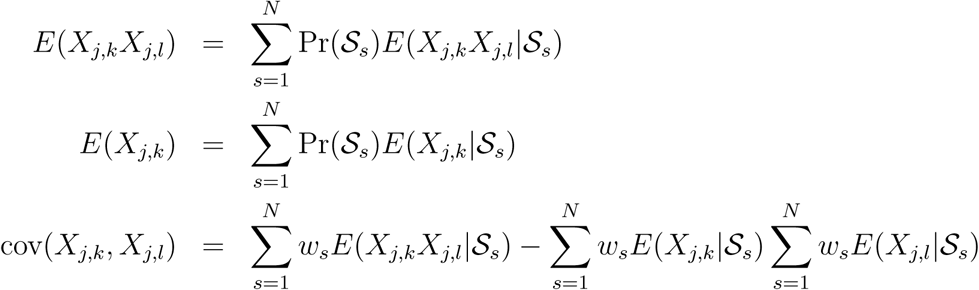

with family weights 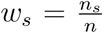 and 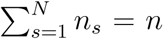. The aim is now to derive an expression that depends on already known terms. For instance, using

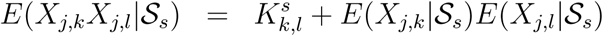

yields

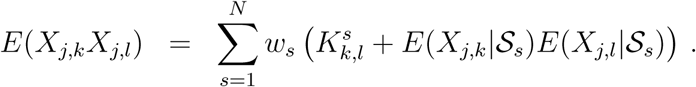

We exploit the separation into independently inherited maternal and paternal SNP alleles: *X*_*j,k*_ = *X*_*j,k,s*_ + *X*_*j,k,d*_, where *X*_*j,k,s*_ and *X*_*j,k,d*_ take a value of 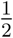 if the A allele was inherited but 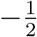 otherwise. Then

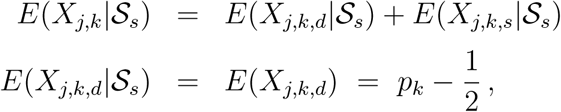

where *p*_*k*_ denotes the maternal allele frequency at SNP *k*. Furthermore,

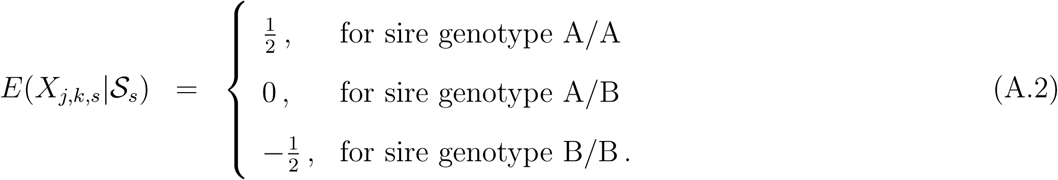

Putting it all together,

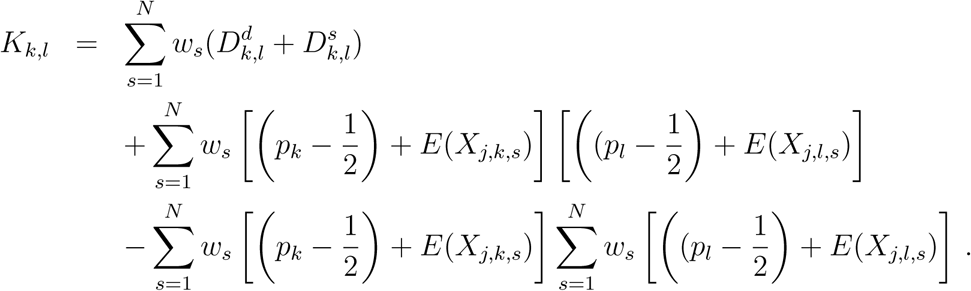

This can be reduced to

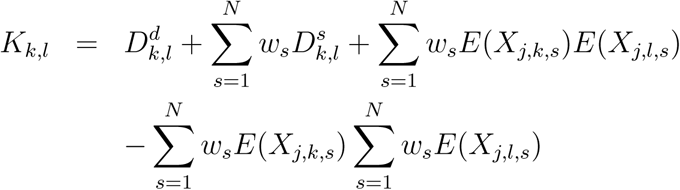

and evaluated using the sire-specific terms in (A.1) and (A.2).

Now the variance of genotype codes at SNP *k* is derived explicitly – it also serves as a scaling term in a regression model for association analysis. The second moment of the paternally inherited SNP allele is constant 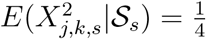 for all *s*. Hence

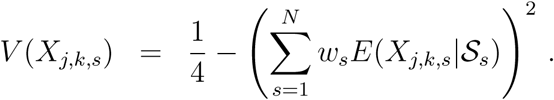

Then, the variance at SNP *k* is

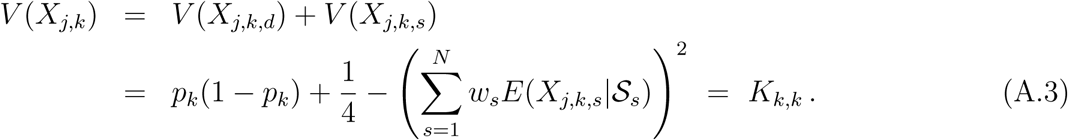

Finally, the correlation matrix 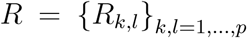 is calculated by scaling the entries correspondingly,

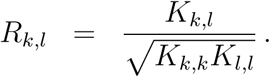

Note that the covariance based on non-centered genotype codes (as derived above) is identical to the one based on centered genotype codes (as used in Section Material and methods). Centering is used to study within-family genetic effects, and it allows the direct estimation of allele substitution effects (Abecasis *et al*., 2000).

### D Supplemental Figures

**Figure D.1.**
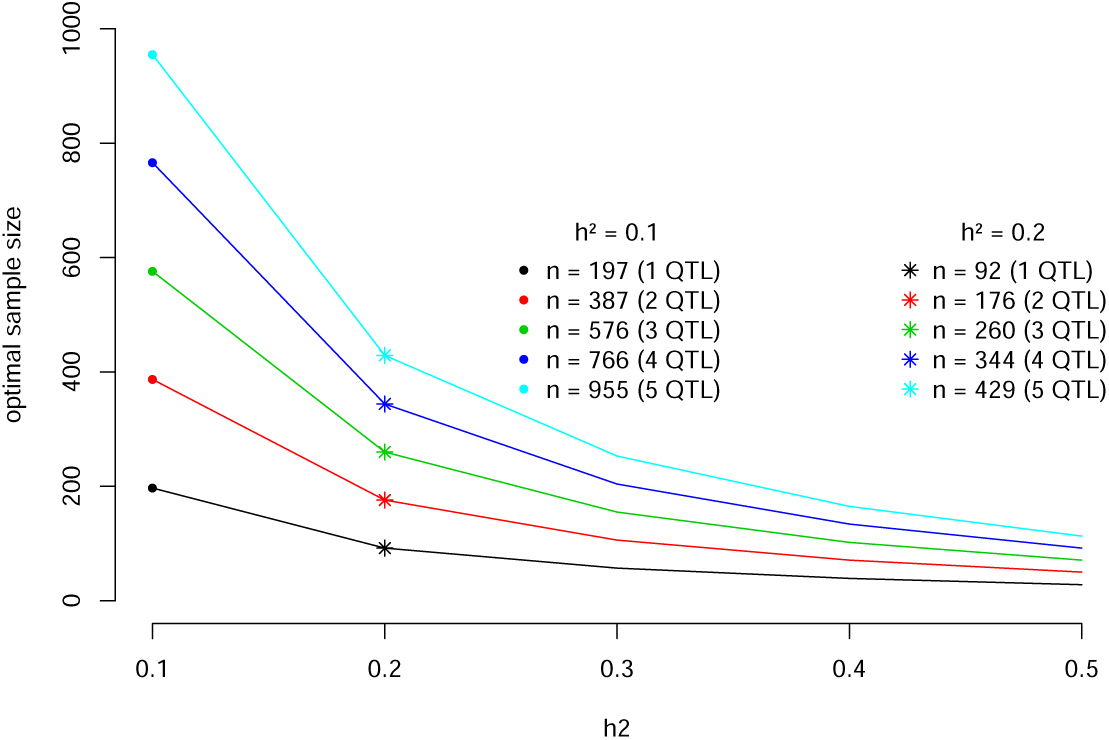
Optimal sample size estimated from the single-SNP model and depending on heritability. Pointwise type-I error was corrected using the simpleℳ method.

**Figure D.2.**
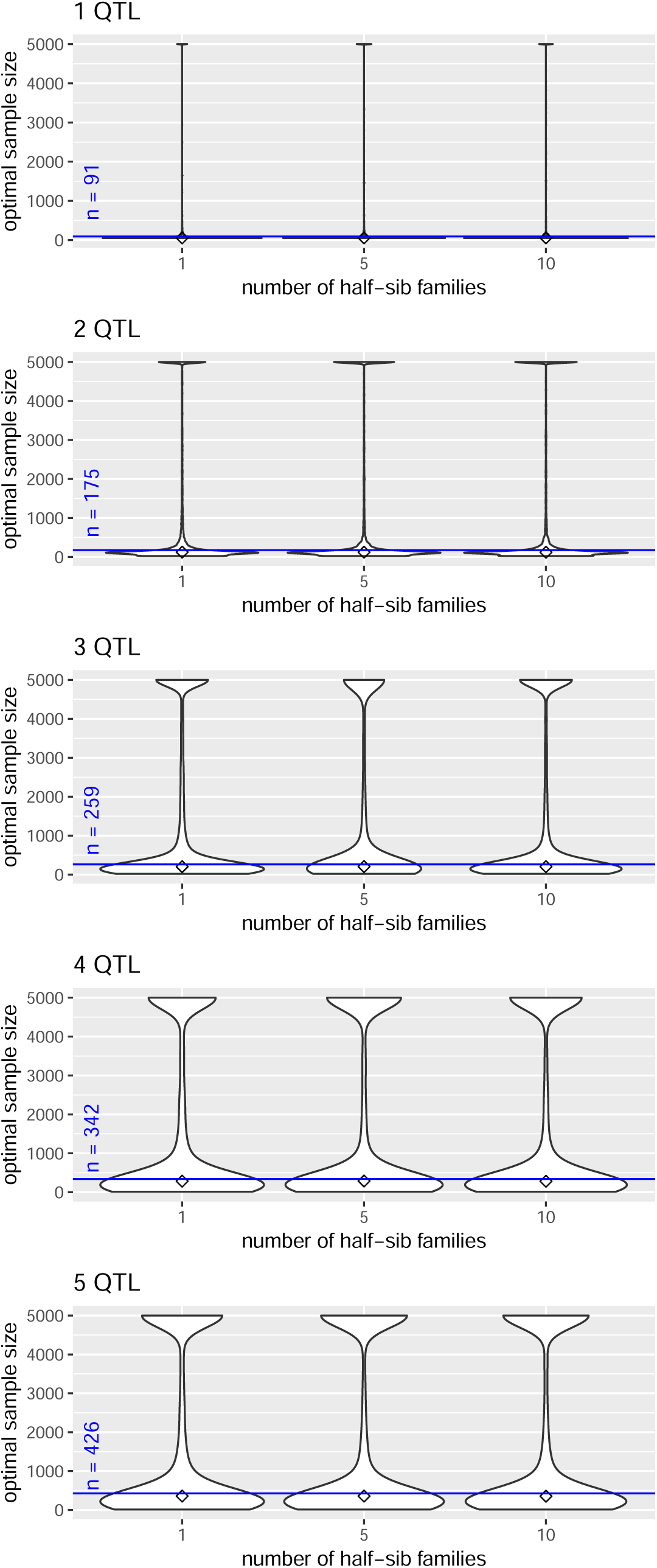
Distribution of optimal sample size. Violinplot of *n*_opt_ vs. number of half-sib families for different numbers of QTL signals in a multi-SNP model. The parent generation was simulated 100 times and 100 random draws of positions of QTL signals were analyzed in each run, *h*^2^ = 0.2. The diamond indicates the median of *n*_opt_ and the blue line marks the results based on a single-SNP model.

**Figure D.3.**
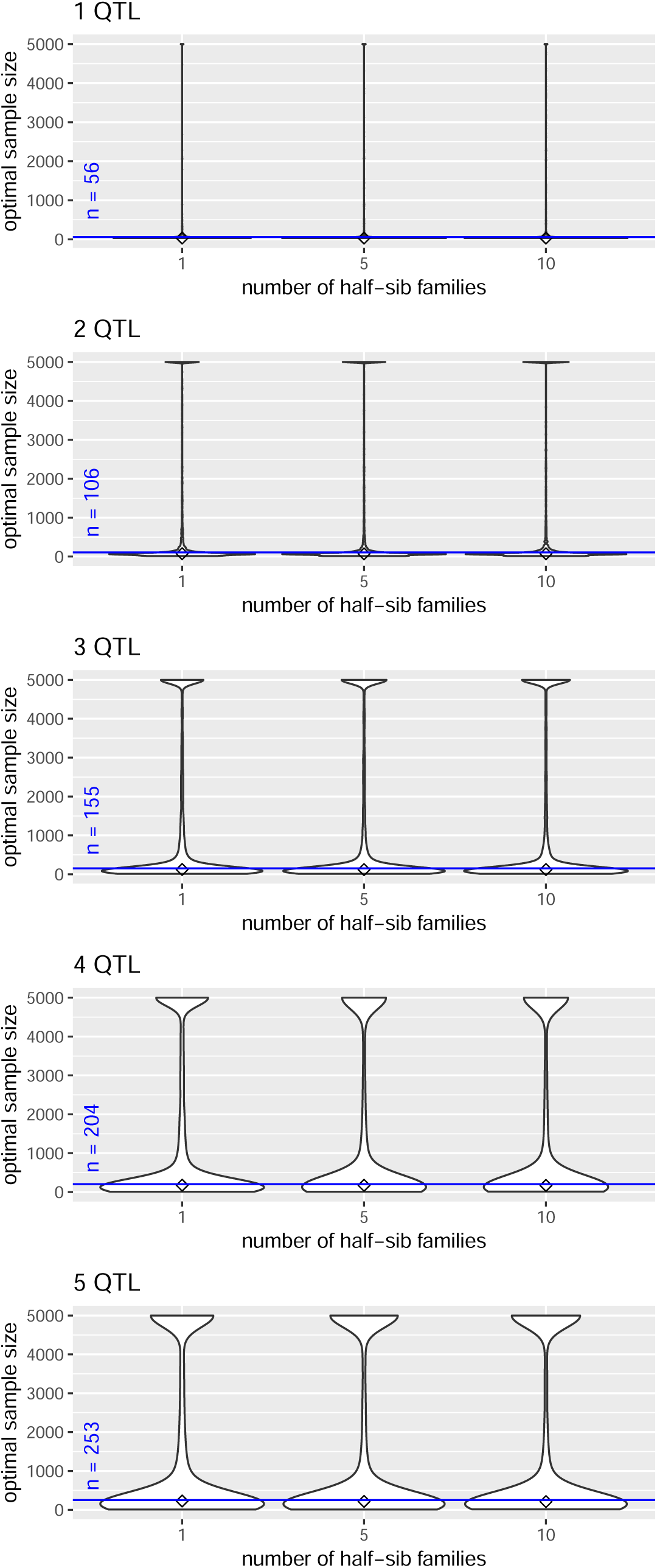
Distribution of optimal sample size. Violinplot of *n*_opt_ vs. number of half-sib families for different numbers of QTL signals in a multi-SNP model. The parent generation was simulated 100 times and 100 random draws of positions of QTL signals were analyzed in each run, *h*^2^ = 0.3. The diamond indicates the median of *n*_opt_ and the blue line marks the results based on a single-SNP model.

**Figure D.4.**
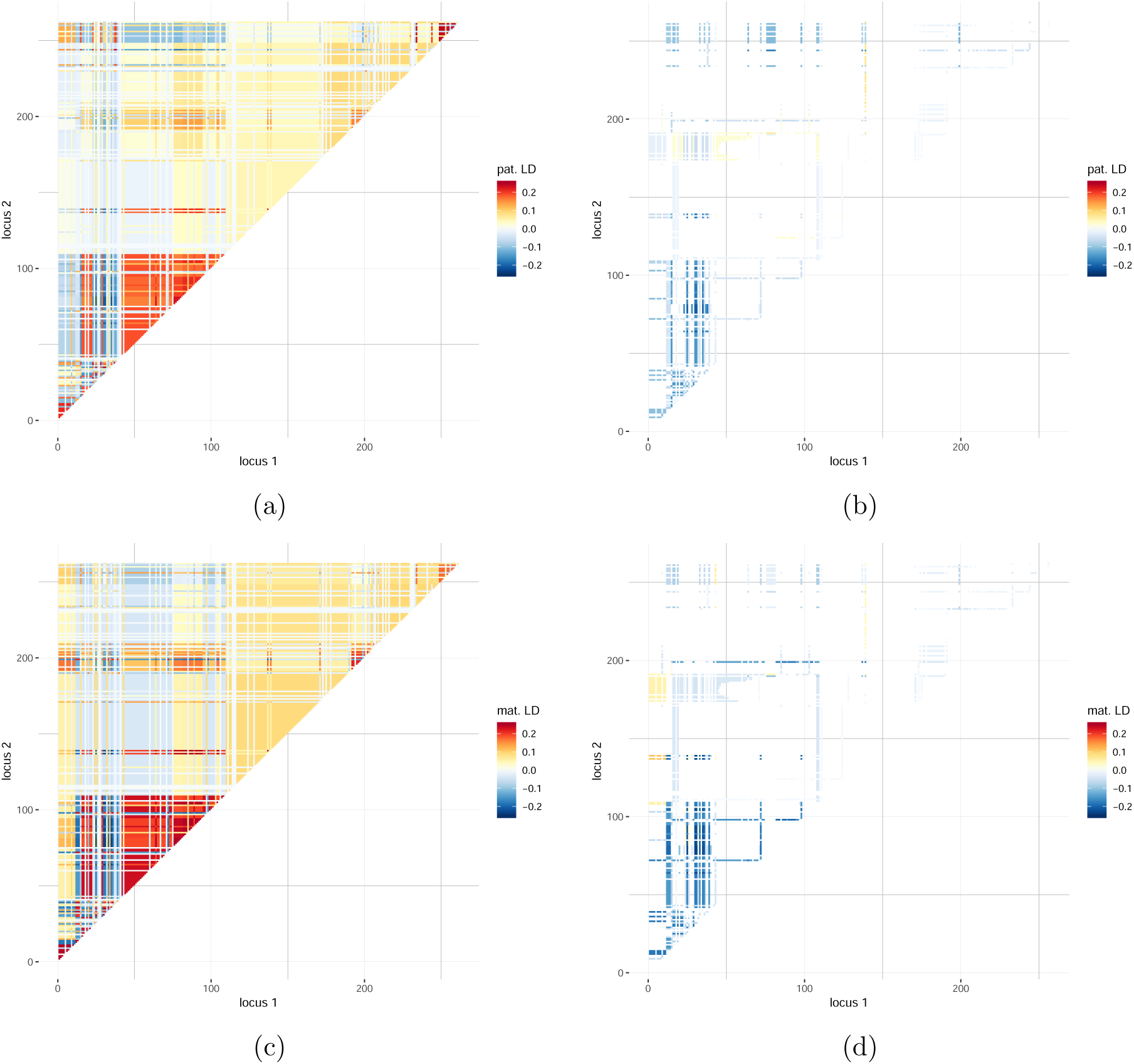
Separation of dependence between SNPs in a single simulated data set with *N* = 10 sires. (a) Paternal covariance, (b) entries selected from paternal covariance which belong to 10% highest sample size (*n*_opt_ ≥ 864), (c) maternal covariance, (d) entries selected from maternal covariance which belong to 10% highest sample size. All possible SNP pairs were evaluated to detect two QTL signals (*h*^2^ = 0.1).

**Figure D.5.**
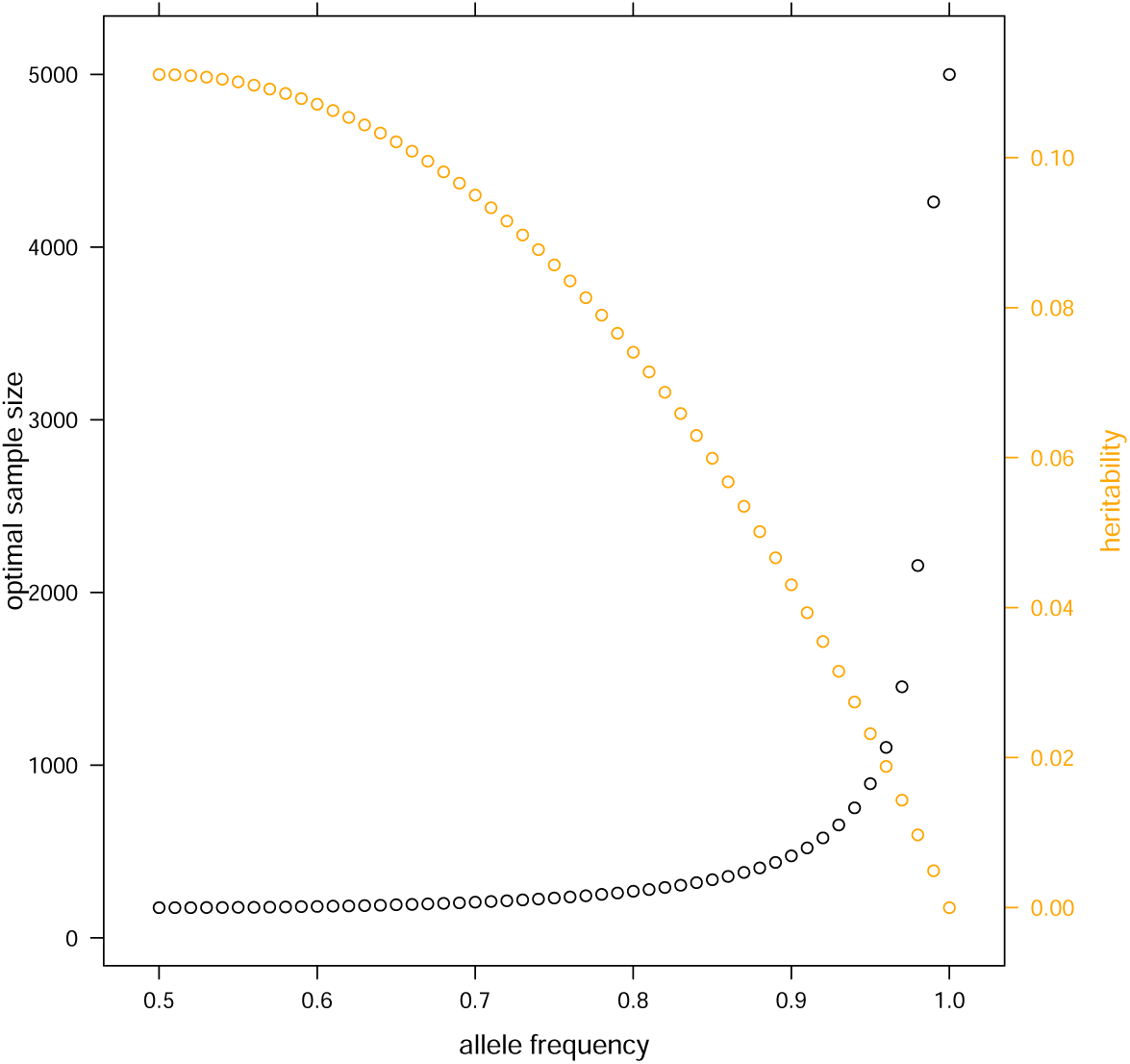
Dependence of optimal sample size on major allele frequency (*p*). The relative effect size on the observed genotype level was fixed at 0.5 and multiplied by 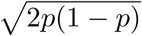. Optimal sample size was estimated based on a single-SNP model.

